# Generation of inheritable A-to-G transitions using adenine base editing and NG-PAM Cas9 in *Arabidopsis thaliana*

**DOI:** 10.1101/2022.06.07.495175

**Authors:** Yi Yun Tan, Yin Yin Liew, Baptiste Castel, Yizhong Zhang, Sang-Tae Kim, Eunyoung Chae

## Abstract

CRISPR/Cas9 technology is an important tool for functional genomics and crop improvement. It can be used to generate mutations at precise positions in the genome. Base editors consist of deaminase components and Cas9 to specify the type of mutation, such as C-to-T (cytosine base editors) or A-to-G (adenine base editors) transition. Available adenine base editor vectors usually make use of canonical Cas9, which limits their use to 5’-NGG-3’ containing targets. We combined a relaxed variant of SpCas9 that uses 5’-NG-3’ containing targets with the adenine base editor containing TadA7.10 or TadA8e to make a set of vectors. By using a phenotype-based screen, we found that our vectors efficiently induce A-to-G somatic mutations in a 5’-NGG-3’ PAM context in *Arabidopsis thaliana* up to 81% efficiency. Such mutations are inheritable at the homozygous stage in T_2_. Among tested vectors, *pECNUS4* (Addgene #184887), which carries TadA8e, showed highest efficiency at generating a stable A-to-G transition in a 5’-NGH-3’ PAM context in the gene *DM3*. Using this vector, we were able to recreate a naturally occurring allele of *DM3* in two generations without the transgene. *pECNUS4* is a new component of the CRISPR toolbox to be used for introducing desired adenine base transitions with an expanded target window for functional genomic research and trait improvement.

## INTRODUCTION

Genome editing has presented itself as a revolutionary tool for engineering novel, desirable traits for crop improvement. The Clustered Regularly Interspaced Short Palindromic Repeat/CRISPR-associated protein 9 (CRISPR/Cas9) system rapidly gained popularity as a powerful tool for genome editing due to its simplicity and versatility (Jinek *et al*., 2012; Razzaq *et al*., 2019). Recapitulating hundreds of years of domestication in tomato was made possible in a matter of months with the CRISPR technology by editing a handful of agronomically important loci in a wild relative (Zsögön *et al*., 2018). Target specificity for the CRISPR/Cas9 system only requires a single guide RNA (gRNA) and the nuclease Cas9. DNA target sites require a protospacer adjacent motif (PAM) next to the target site at the 3’ end, recognised by the Cas9 protein (Mojica *et al*., 2009). Cas9 cleaves the DNA at a target locus complementary to the gRNA to produce double-stranded breaks (DSBs), which are then repaired by homology directed repair (HDR) or non-homologous end-joining (NHEJ) pathway to lead to desired sequence alterations (Yin *et al*., 2017).

Single nucleotide polymorphisms (SNPs) are often determining factors in expression of favourable traits (Mojica *et al*., 2009; Li *et al*., 2017). The conventional CRISPR/Cas9 application causes random small insertions, deletions or base replacement at the target. The development of base editors, adenine base editors (ABEs) and cytosine base editors (CBEs), has allowed CRISPR/Cas9 technology to expand its scope from molecular scissors to precise nucleotide converters. ABE and CBE have been applied to produce targeted, point mutations in numerous organisms via deaminase components fused to a Cas9, when directed to a target position by a single gRNA (Komor *et al*., 2016; Gaudelli *et al*., 2017). Undesirable double-stranded breaks are avoided with the use of Cas9 mutated in one or both of its two nuclease sites. Through an adenosine deaminase enzyme tethered to a Cas9 nickase, ABEs introduce a base change of adenine to inosine, which is fixed to guanine by DNA polymerases (Gaudelli *et al*., 2017).

Compared to the highly efficient CBEs, ABEs appear to produce relatively lower mutation rates in plants (Kang *et al*., 2018; Hua *et al*., 2019; Molla and Yang, 2019; Negishi *et al*., 2019; Zeng *et al*., 2020). Recently, an engineered adenosine deaminase, TadA8e, has been reported to exhibit up to ∼1100-fold faster DNA deamination kinetics than the first generation deaminase TadA7.10, with eight amino acid substitutions (Richter *et al*., 2020; Yan *et al*., 2021). Substantially improved base-editing efficiencies were observed with TadA8e both in mammalian cells and in rice (*Oryza sativa*) (Huang *et al*., 2021; Hua *et al*., 2022; *et al*., 2022). TadA9 with one additional mutation compared to TadA8e was also shown to have increased base-editing efficiency in rice (Yan *et al*., 2021). Efficient adenine base editing has been reported in several dicot plants including allotetraploid cotton (Niu *et al*., 2021; Wang *et al*., 2021, 2022). Applications of ABEs have been described in *Arabidopsis thaliana* with limited editing efficiencies even with TadA8e (Mao *et al*., 2021).

5’-NGG-3’ PAM requirement by the most widely used Cas9 allele (from *Streptococcus pyogenes* SpCas9) restricts the number of targets on the genome. Multiple Cas9 variants were developed via rational engineering and directed evolution to have altered PAM specificities, including SpCas9-NG (Walton *et al*., 2020, Nishimasu *et al*., 2018, Kleinstiver *et al*., 2015, Slaymaker and Gaudelli, 2021). Such variants have been shown to work with relatively high base-editing efficiencies in rice (Li *et al*., 2021; Ren *et al*., 2021; Wei *et al*., 2021; Yan *et al*., 2021; Tan *et al*., 2022). The near-PAMless SpRY-TadA8e seemed to be highly efficient in mediating A-to-G base editing, albeit with added considerations for experimental design in regard to T-DNA self-editing and off-target effects.

In this study, we developed four vectors for adenine base editing in *A. thaliana* containing TadA7.10 or TadA8e as a deaminase, paired with a plant-codon optimised Cas9 variant SpCas9-NG that recognizes 5’-NG-3’ PAM sites, with or without an intron (Nishimasu *et al*., 2018; Castel *et al*., 2019; Zhong *et al*., 2019). We first assessed the base editing capabilities of our ABE vectors for 5’-NGG-3’ PAM context using a previously described albino reporter system (Kang *et al*., 2018). Second, we tested the base editing capabilities of the four ABE vectors in the 5’-NGH-3’ PAM context. We particularly aimed to generate naturally occurring alleles of an immune gene *Dangerous Mix 3* (*DM3*) in *A. thaliana* (Chae *et al*., 2014). One vector, *pECNUS4*, has an overall high efficiency and was used to produce a T-DNA-free homozygous base-edited mutant in two generations for *PDS3* (with 5’-NGG-3’ PAM) and *DM3* (with 5’-NGH-3’ PAM). The ABE vector carrying TadA8e, *pECNUS4*, produced T_1_ lines with up to 81% mutation rate detected at somatic level in the *PDS3* 5’-NGG-3’ context, and 40% in the *DM3* 5’-NGC-3’ context. A stable, transgene-free homozygous mutant was found upon surveying nine of the Cas9-free T_2_ progeny of the 40% edited T_1_ line. In all, we demonstrated stable A-to-G base changes in T-DNA-free plants in T_2_ generation for targets with canonical 5’-NGG-3’ PAM sites and non-canonical 5’-NGH-3’ PAM sites in *A. thaliana*.

## RESULTS

### Efficiency of somatic A-to-G editing with 5’-NGG-3’ PAM and its heritability

We generated four vector constructs, namely *pECNUS1, pECNUS2, pECNUS3*, and *pECNUS4*, with one of the two adenosine deaminase variants, TadA7.10 or TadA8e, and one of the two SpCas9-NG variants (Figure 1A) (Gaudelli *et al*., 2017; Richter *et al*., 2020). One *Cas9* allele contains a *potato intron IV* and the other does not (Castel *et al*., 2019; Zhong *et al*., 2019). The constructs contain an *RPS5A* promoter to drive expression of Cas9 fused to the respective deaminase and an RNA polymerase III-expression cassette for gRNA expression. A 20-nt double stranded spacer sequence flanked by ATTG-GTTT overhangs can be inserted by digestion-ligation in a single reaction. After this step, the construct can be expressed in *A. thaliana* by *Agrobacterium*-mediated transformation (Figure 1B). It contains a FASTRED selection marker, which renders the seeds fluorescent and easily selectable under a microscope, thus enabling selection of Cas9-free lines after the generation of the desired editing event. We assessed the editing activity at somatic level in T_1_, using phenotyping and amplicon deep sequencing (Figure 1B), and the inheritability of the events in T_2_ using phenotyping and Sanger sequencing.

**Figure 1.**
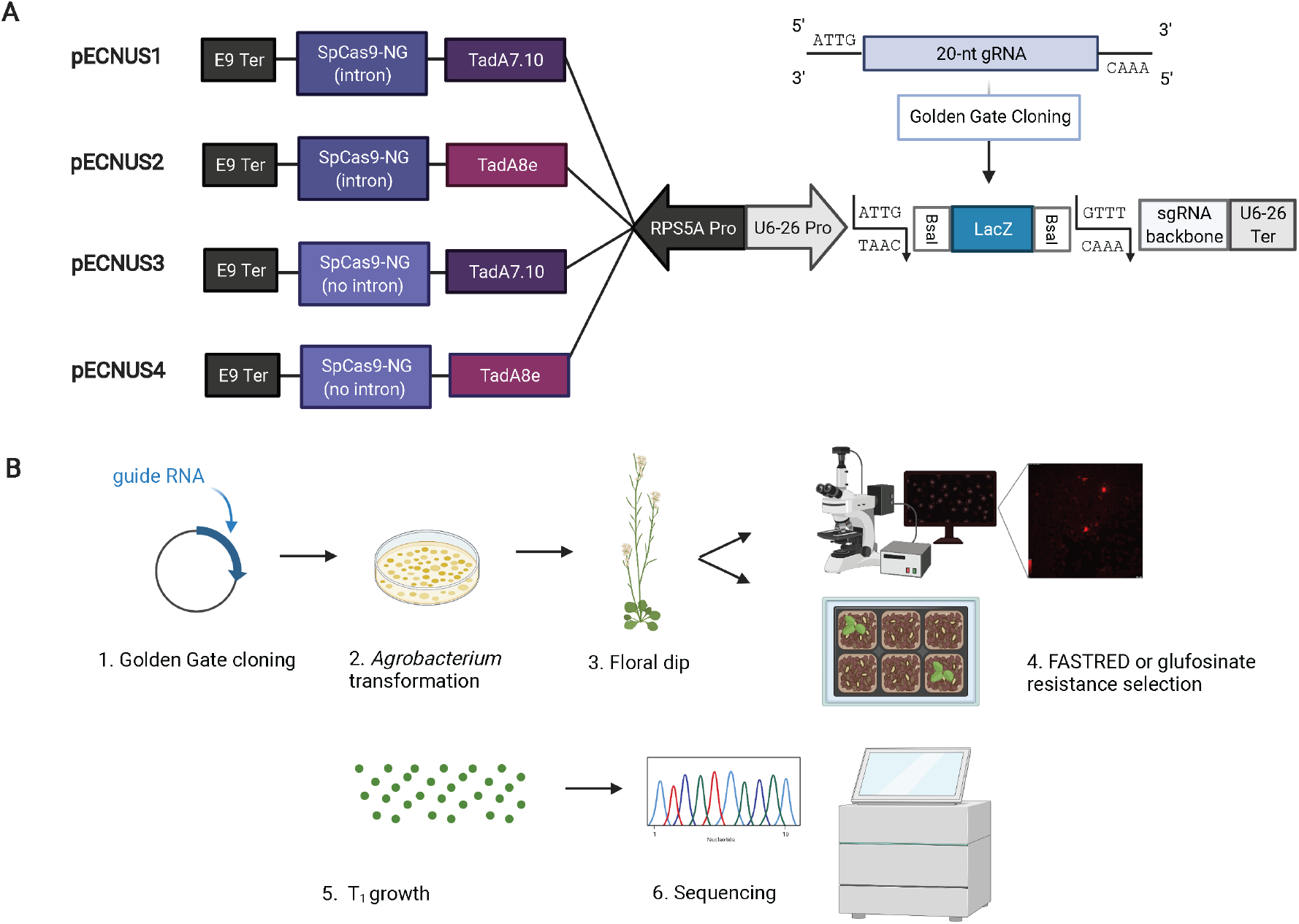
Workflow for establishing heritable ABE in *A. thaliana*. **A**. Schematic diagram of ABE vector constructs *pECNUS1, pECNUS2, pECNUS3*, and *pECNUS4* containing two adenosine deaminase variants: TadA7.10 or TadA8e, and one of two SpCas9-NG sequence variants. The LacZ selection marker is flanked by *Bsa*I restriction sites and ATTG/GTTT overhangs. Insertion of a 20-bp gRNA target sequence with complementary overhangs can be performed via a Golden Gate reaction into the ABE vectors, replacing the LacZ selection marker. Each vector construct contains both FASTRED cassette and glufosinate resistance as plant selectable markers. **B**. Schematics of workflow for generating ABE transgenic *A. thaliana* plants.

First, we tested the efficiency of the constructs carrying the TadA8e variant, which are *pECNUS2* and *pECNUS4*, using an albino reporter system previously reported (Kang *et al*., 2018). It relies on targeted base changes at a splice-acceptor site of *PDS3*, for which a gRNA can be designed with 5’-NGG-3’ PAM. A successful ABE event causes mis-splicing of *PDS3* resulting in loss-of-function and an albino phenotype (Figure 2A). We cloned this gRNA in *pECNUS2* and *pECNUS4* and assessed the editing efficiency in *A. thaliana*. Guided by albino phenotype, we were able to determine which individuals were affected and thereby compare the ABE efficiency of different constructs in T_1_. We then confirmed the editing efficiency by amplicon deep sequencing using Illumina iSeq 100. For both constructs *pECNUS2* and *pECNUS4*, we observed albino phenotypes among the T_1_ individuals carrying either of the constructs. Some T_1_ plants showed albino symptoms in rosette leaves, cauline leaves and/or floral heads. We also observed milder manifestations with pale green sectors (Figure 2A). For one T_1_ plant manifesting mosaic phenotype, we collected DNA from various white and pale green parts and measured the A-to-G conversion rate by Sanger sequencing and deep sequencing (Figures 2A and S1). At an early stage, we found that the A-to-G conversion rate was high, up to 73%, in a white sector, while the rate ranges from 11% to 15% in the green sectors (Figure S1). At a later stage, we found that for the white sectors, the conversion rates were greater than 50%, while 40% was seen in pale green tissue (Figure 2A). These results show that independent ABE events can occur in a mosaic pattern in T_1_.

**Figure 2.**
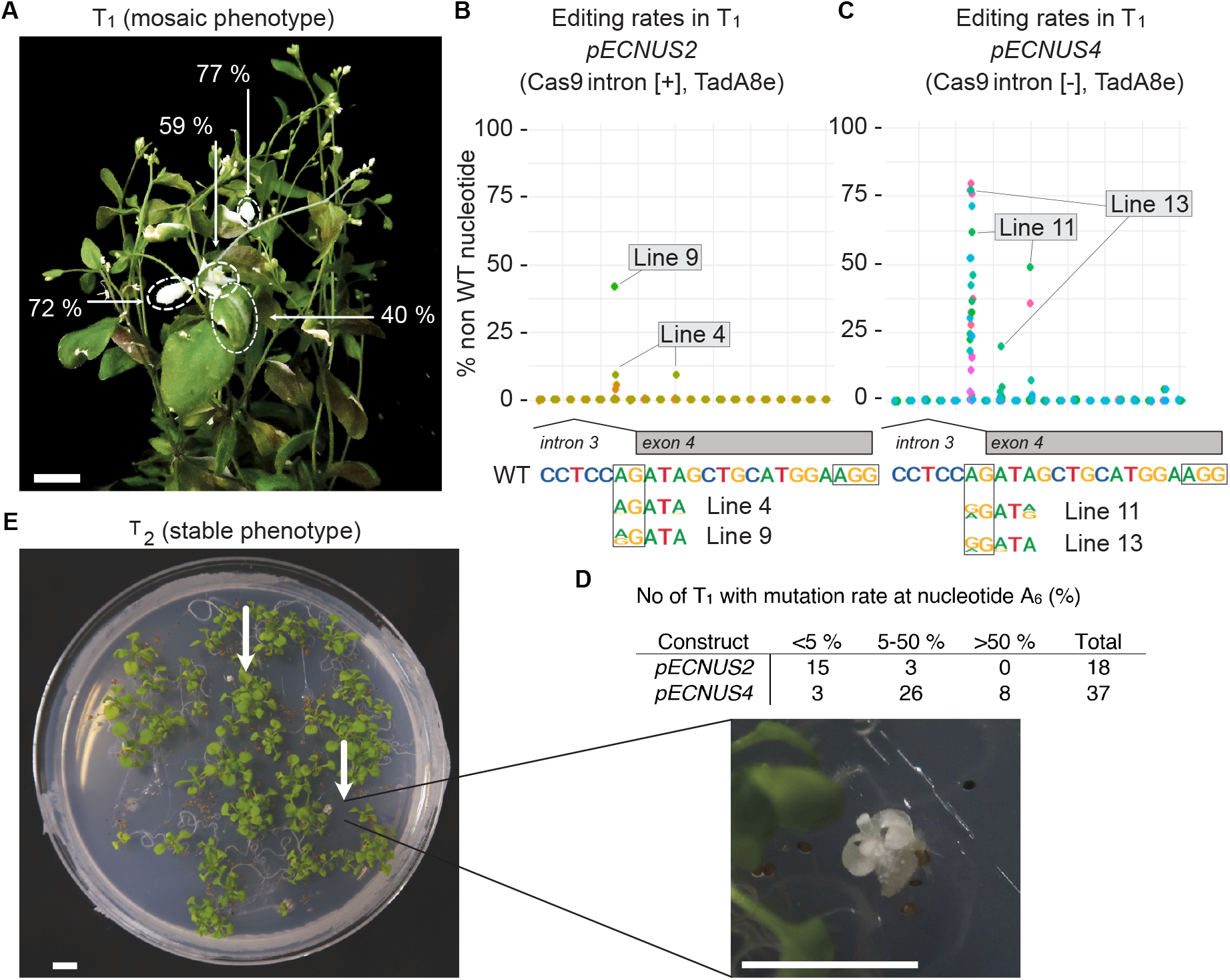
Adenine base editing at *PDS3* target site with 5’-NGG-3’ PAM. **A**. Mosaic phenotype in a T_1_ plant expressing *pECNUS4* with gRNA targeting *PDS3*. The plant is overall green with mosaic occurrences of white. DNA was collected from the area indicated with white dashed line and analysed by deep sequencing. Numbers indicate the percentage of ABE-induced editing events. Bar indicates 1 cm. **B.C**. Efficiency of *pECNUS2* and *pECNUS4* at the *PDS3* target in T_1_, expressed in percentage of base conversion at each site of the gRNA target. Each colour represents one of 18 (B) or 32 (C) independent T_1_. A diagram of the intron / exon limit and the WT bases are indicated below the plot. The ‘AG’ splicing site is framed with black line. The sequence of samples from Lines 4 and 9 is indicated below the WT sequence for *pECNUS2*. The sequence of samples from Lines 11 and 13 is indicated below the WT sequence for *pECNUS4*. The relative size of the bases represents their respective ratios within the total sample reads. Only ‘AGTA’ is represented as the rest of the sequence is not targeted by ABE (no A and out of the predicted editing frame). **D**. Number of T_1_ with mutation rates at nucleotide A position 6 (PAM being at position 20-23). **E**. Stable albino phenotype in T_2_ plants. The seeds from a single independent T_1_ were sown on a MS½ petri dish. Two plants show a complete albino phenotype, indicating complete absence of PDS3 activity. Bar indicates 1 cm.

We used deep sequencing to determine the base editing efficiency at each site, by calculating the percentage of amplicon reads that had none of the wild-type bases at the target site (Figure 2B, 2C). *pECNUS4* showed high adenine base efficiency with eight T_1_ individuals out of 37 having more than 50% base-conversion rates, reaching up to 81%, at the preferred target site A at position 6 (A_6_) (Figure 2C, 2D). This site falls within the expected base targeting window ranging position 4 to 9, relative to PAM at position 21-23. We also detected appreciable efficiency with *pECNUS2*, albeit lower than that of *pECNUS4* in its distribution of edited T_1_ plants (Figure 2B, 2D). Nonetheless, both constructs also resulted in double editing events at positions ranging from 6 to 10 (Line 4 in Figure 2B; Line 11 and Line 13 in Figure 2C). To test whether mutations could be inherited, we screened for full albino plants in T_2_ (Figure 2E). From a line carrying *pECNUS4*, two dwarfed albino seedlings were observed out of a pool of 82 T_2_ lines, indicative of homozygous mutation. Genotyping for Cas9 shows that Cas9 T-DNA was absent from the genomic DNA of the two albino seedlings (Figure S2), confirming heritability. Taken together, our phenotype-guided assessment of editing efficiency revealed that *pECNUS4* could generate substantial editing events for a target with 5’-NGG-3’ PAM at somatic level in mosaics in T_1_, which can be inherited as a stable homozygous mutation in T_2_.

### Inheritable A-to-G editing events at 5’-NGH-3’ PAM target

We further tested the efficiencies of the ABE constructs on target sites with non-canonical 5’-NG-3’ PAMs in *A. thaliana. DM3* was chosen as a target gene given that a natural allele could trigger immune responses in a certain genetic background of *A. thaliana* (Chae *et al*., 2014). When we designed non-synonymous mutations in *DM3* (Q171 and T359) (Figure 3A), we took into consideration that the editing events generally occur between positions 4 and 9 in the gRNA (Gaudelli *et al*., 2017). The gRNA targeting T359 presents a 5’-NGC-3’ PAM, and its editing frame 4-9 contains three As, one in the codon GAT (D358) and two in the codon ACA (T359) (Figure S3A). A-to-G conversion could result in GGT (D358G) and/or in GCA/GCG (T359A), ACG (T, synonymous), while our desired mutation was T359A. The gRNA targeting Q171 presents a 5’-NGA-3’ PAM, and its editing frame 4-9 contains two As, both in the codon CAA (Q) (Figure S4). A-to-G conversion could result in CGA/CGG (R), or CAG (Q, synonymous), while our desired mutation was Q171R.

**Figure 3.**
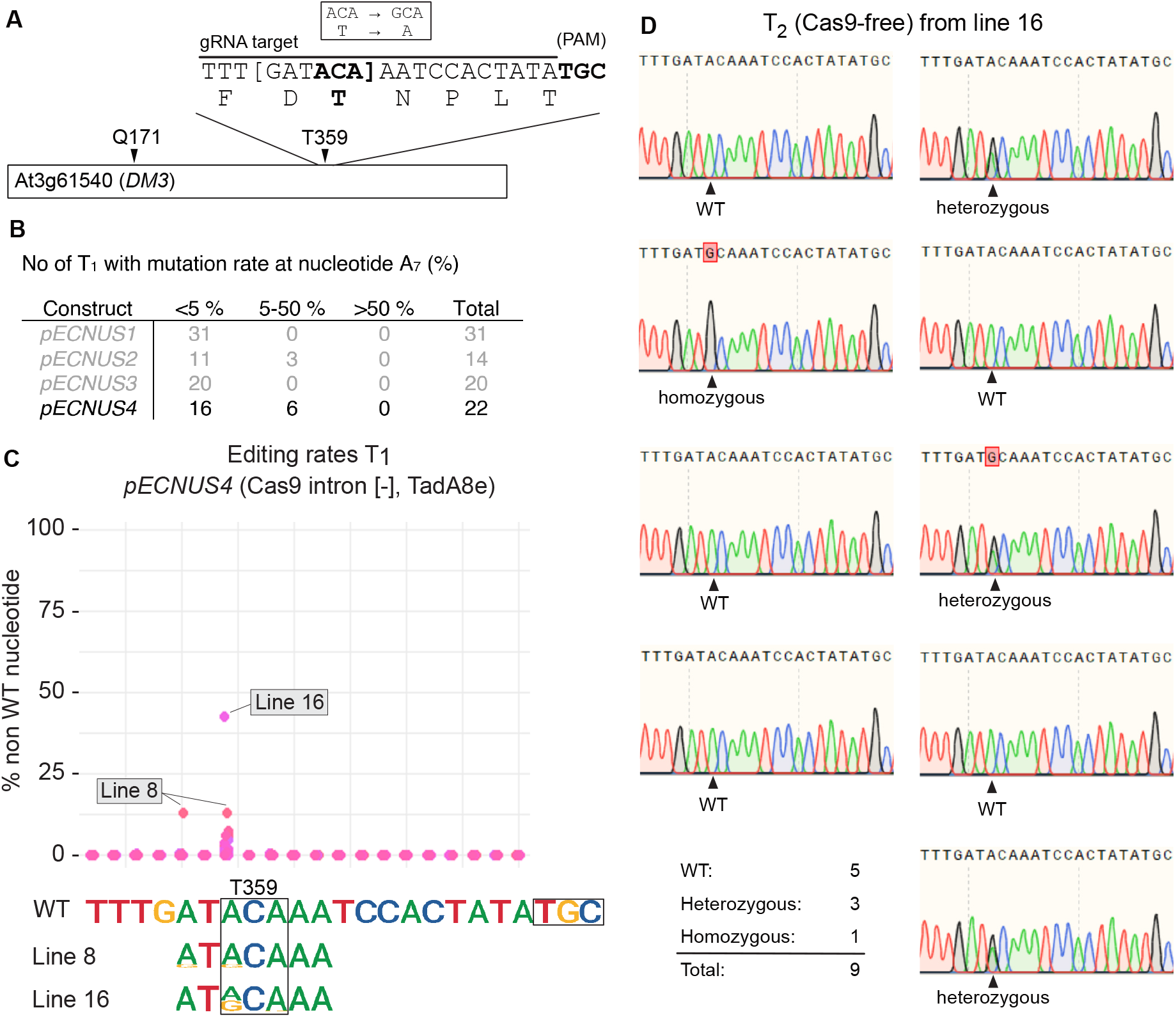
Adenine base editing at *DM3_T359* target site with 5’-NGH-3’ PAM. **A**. Diagram representing the T359 target in *DM3*. Encoded amino acids are indicated below the corresponding codons below the gRNA target sequence. The predicted editing frame 4-9 is delimited by brackets. The Q171 target is presented in Figure S4. **B**. Number of T_1_ with mutation rate at nucleotide A position 6 (PAM being at position 20-23) for each of the four vectors. Details of *pECNUS1, pECNUS2* and *pECNUS3* are presented in Figure S3. **C**. Efficiency of *pECNUS4* at the *DM3_T359* target in T_1_, expressed in percentage of base conversion at each site of the gRNA target. Each colour represents one of 22 independent T_1_. The WT bases are indicated below the plot. The ‘ACA’ coding encoding T is framed with black line. The sequence of samples from Lines 8 and 16 is indicated below the WT sequence. The relative size of the bases represents their respective ratios within the total sample reads. Only ‘ATACAAA’ is represented as the rest of the sequence is not targeted by ABE (no A or out of the predicted editing frame). **D**. Sanger sequencing of nine individual T_2_, T-DNA-free progeny (selected against FASTRED) of the single independent T_1_ Line 16. Five lines indicate a clear single peak for WT residue A, one indicates a clear single peak for edited residue G (homozygous) and three indicate a dual peak (heterozygous).

First, we tested the gRNA targeting T359 with 5’-NGC-’3 PAM in all four vectors. The ABE constructs containing TadA8e vectors *pECNUS2* and *pECNUS4* induced higher base-conversion rates than the TadA7.10 vectors *pECNUS1* and *pECNUS3* (Figures 3B and S3). Deep sequencing identified three and six T_1_ individuals with editing rates of 5-50 % at target nucleotide A_7_ for *pECNUS2* and *pECNUS4*, respectively. Of note, Line 16 carrying *pECNUS4* was shown to have a 40% base-editing event at the target nucleotide A_7_ in T_1_, inducing the desired T359A mutation only. No significant base editing events were observed for the T_1_ lines transformed with *pECNUS1* and *pECNUS3* (Figure S3). We tested our most efficient construct, *pECNUS4*, with the gRNA targeting Q171 with 5’-NGA-3’, but we did not obtain significant base editing events (Figure S4).

To observe heritable ABE-induced mutations, we screened for Cas9-free T_2_ progeny of Line 16 by selecting for FASTRED-negative seeds under fluorescent microscopy. Since these lines are Cas9-free, they cannot contain *de novo* editing events; all the mutations observed are necessarily inherited from the parent. From nine Cas9-free progeny, we identified five wild-type individuals and three heterozygous and one homozygous offspring carrying the mutation observed in T_1_ (Figure 3D). We reason that deviation from the 1:2:1 segregation is due to heterogeneous editing events that occurred on inflorescence tissues. This result confirms that the vector *pECNUS4* could generate a T-DNA-free homozygous A-to-G base-edited mutant in two generations in a 5’-NGH-3’ context.

## DISCUSSION

In this study we present a vector, *pECNUS4*, which can generate A-to-G base editing at both canonical 5’-NGG-3’ PAM and non-canonical 5’-NGH-3’ PAM targets in *A. thaliana*. We used deep sequencing in T_1_ to assess the overall efficiency of four constructs. In T_1_, we observed somatic events occurring in mosaic patterns in the plant. We found heterogeneous results, from no activity to 81% edited reads in independent transgenic plants (Figure 2C). Heterogeneity occurs both between plants (Figures 2B, 2C and 3C) and within a single plant (Figure 2A). We also found varying results for different targets. For instance, with the same vector *pECNUS4*, we found 8 out of 37 samples with more than 50% base-conversion rates for *PDS3* target with 5’-NGG-’3 PAM (Figure 2D); while all samples showed below 50% edition for the *DM3_T359* target with 5’-NGC-’3 PAM. We did not detect any activity for the *DM3_Q171* target with 5’-NGA-’3 PAM (Figure S4). It is worth noting that the sampling method used for *PDS3* is biased and assisted by albino phenotype. The higher rates detected for *PDS3* suggests that actual base editing events could have been underestimated when not guided by phenotypic cues. Nonetheless, it is well known that gRNAs have varying efficiencies, although the reasons are not clear. The software CRISPRon (Xiang *et al*., 2021), recently developed and based on deep learning, predicts an on-target activity of 48% for our gRNA targeting *PDS3*, while it predicts 19% and 21% for the gRNA targeting *DM3* at T359 and Q171 targets respectively (assuming an 5’-NGG-3’ PAM). Another reason for different editing efficiencies could be due to the use of non-canonical PAM to target T359 and Q171. Indeed, engineered Cas9 alleles with relaxed PAM generally show a higher activity at 5’-NGG-3’ PAM than the non-canonical ones (Chatterjee, Jakimo and Jacobson, 2018; Zhong *et al*., 2019; Ren *et al*., 2021).

Interestingly, even in the absence of albino phenotype, 15% base-conversion rate was still observed in T_1_ (Figure S1), indicating that basal levels of base editing could occur even if the expected phenotype does not appear. In our non-phenotype guided assessment, we observed below 50% efficiency in most T_1_ samples, whereas the line with 40% somatic efficiency was sufficient to produce inheritable mutation. Hence, genotyping is essential to screen for ABE activity in T_1_ to screen for heritable mutations. In this study, we used deep sequencing to measure and report precisely the efficiency of our constructs. To obtain at least one line with high mutation rates, we recommend the users to test at least ten independent transformants for 5’-NGG-3’ PAM and at least 40 for 5’-NGH-3’ PAM. When possible, it is preferable to choose a 5’-NGG-3’ PAM target for the use of *pECNUS4*.

ABE events are expected to occur in A at position 4 to 9 in the target, considering the PAM at position 21 to 23 (Gaudelli *et al*., 2017). Indeed, we observed the highest editing rates at the position A_6_ for *PDS3* (Figure 2BC) and at A_7_ in *DM3_T359* (Figure 3C). We could also see activity at As positioned at 5 (Figure 3C), 8 and 10 (Figure 2BC). It indicates that our construct *pECNUS4* can induce A-to-G mutations in an editing frame of at least 5 to 10. We also assessed off-target effects in successfully base-edited T_2_ lines from Line 16 at a predicted off-target site with two mismatches (Figure S5) and did not find any signs of editing (Figure S5). While more extensive off-target screens could be conducted, ABE was reported not to have significant off-target effects (Zuo et al., 2019; Jin et al., 2019).

The addition of an intron in Cas9, originally proposed to avoid expression in bacteria during cloning (Li *et al*., 2013), was expected to enhance the overall CRISPR activity, likely by allowing more stable expression (Castel *et al*., 2019; Grützner *et al*., 2021). Surprisingly, we observed the opposite here. With the same gRNA, the construct *pECNUS4* without intron induced more editing events than the construct *pECNUS2* with intron (Figure 2D and 3B). It is at present unclear why the presence of an intron did not improve Cas9 expression for our vectors.

Our objective was to generate a CRISPR construct for A-to-G base editing combined with relaxed PAM requirements to provide a tool to mutate SNPs for functional genomics and trait improvement. Our results present *pECNUS4* as a suitable vector for this purpose that can generate high rates of editing events in primary transformants that can produce progeny carrying the inherited mutation in a transgene-free homozygous state in two generations. Any species amenable for *Agrobacterium*-mediated transformation with adequate transformation efficiency will benefit the use of the presented tool for trait improvement.

## Materials and Methods

### Vector cloning

The vectors were assembled using the Golden Gate modular cloning method for plants (Patron *et al*., 2015). Key components of the binary vectors have been published (Castel *et al*., 2019) and assembled in Level 2 acceptor vectors with kanamycin resistance. Primers flanked with *Bsa*I restriction sites and Golden Gate compatible overhangs (Patron *et al*., 2015) were used to amplify respective components for the Level 0 vector (Table S1 and S2). *A. thaliana* codon-optimised coding sequence of TadA8e was synthesised with flanking *Bsa*I sites (IDT; sg.idtdna.com/) and cloned into respective Level 0 vectors containing Cas9 variants. Domestication to remove a *Bsa*I restriction site from the TadA8e coding sequence was performed with primers as specified (Table S1). The LacZ selection marker was PCR amplified with flanking *Bsa*I sites into Level 1 acceptor vector *pICH41780* (Addgene #48019) to allow blue-white screening for successful gRNA insertion into final ABE vectors. Details about the reaction mix and thermocycler programs used for construct generation are found in Supplemental Methods 1.

### gRNA cloning into the vector

Cloning of final ABE vectors containing gRNA were performed as follows. Forward and reverse oligos were synthesised with respective overhangs by IDT (sg.idtdna.com) to form gRNA with sticky ends, using annealing protocol previously reported (Kim *et al*., 2016). The gRNAs were cloned into ABE vectors with equimolar volumes via a digestion-ligation reaction (diglig) with *Bsa*I enzyme. The diglig mix was transformed into *E. coli* DH5alpha and selected on media with kanamycin. White positive colonies were selected via blue-white screening and plasmid DNA were isolated using EZ-10 Spin Column Plasmid DNA Miniprep Kit (Bio Basic Asia, Singapore). Sequence information of all cloned vectors were confirmed with Sanger sequencing (Bio Basic Asia, Singapore).

### Plant growth and transformation

*A. thaliana* ecotype Col-0 was used in this study. Plant growth was carried out at 23°C under long-day conditions (16 hours of light / 8 hours in dark) in a plant growth room. All constructs were transformed into *A. thaliana* via *Agrobacterium*-mediated floral dip transformation with *Agrobacterium tumefaciens* strain GV3101 (Clough and Bent, 1998). Transformants (T_1_) were selected using a FASTRED marker under fluorescent microscopy. Seeds were sterilised using a seed wash solution (70% ethanol, 0.05% Triton X-100), followed by 95% ethanol. The seeds were germinated on half-strength Murashige and Skoog (MS½) medium following a stratification period of three days in 0.1% (w/v) agar in the dark at 4°C. T_1_ hybrids were grown in chambers at 23°C under long-day conditions with light supplied by Philips GreenPower TLED modules (Philips Lighting GmbH, Germany). After 4 weeks of growth, plants were transferred to soil and grown under long-day conditions at 23°C for flowering.

### Genotyping by sequencing

Cetyltrimethylammonium bromide genomic DNA extraction was performed for samples collected from T_1_ plants (Aboul-Maaty and Oraby, 2019). Target loci were amplified using Taq DNA polymerase (Bio Basic Asia) with appropriate primers (Table S2). Library preparation for targeted deep sequencing was performed via three steps of PCR using site-specific primers (Table S2). First PCR is used to isolate the general region of interest from genomic DNA, and second PCR further defines region for deep sequencing and adds flanking sequences complementary to third PCR primers, which adds flanking indexes to the amplicons. Pooled purification of the PCR amplicons was then performed using Solid Phase Reversible Immobilisation (SPRI) beads (Beckman Coulter, Singapore). Purified PCR amplicons were sequenced via next-generation sequencing (NGS) by Illumina iSeq 100 Sequencing System following manufacturer recommendations. NGS paired-end reads were processed using the ‘fastq-join’ program from the ‘ea-utils’ package (Aronesty, 2013).

NGS output files were further processed using a modified python script (Choi *et al*. 2021). The files of the NGS data generated after processing can be found in Supplemental Dataset 1. Base conversion rates from A-to-G were additionally visualised via BE-Analyzer (Hwang *et al*., 2018). Possible off-target sites were detected via CasOFFinder (Bae *et al*. 2014). PCR products of selected genomic loci were also sequenced by Sanger sequencing (Bio Basic Asia, Singapore).

## Acknowledgments

We thank Donghui Hu and Wei-Yuan Cher for technical support. This work is supported by Singapore National Research Foundation under its Competitive Research Programme (NRF-CRP22-2019-0001). S.T.K was funded by the National Research Foundation of Korea (NRF) grant funded by the Korean government (MSIT) (No. 2022R1A2C1011598). The funders had no role in study design, data collection and analysis, decision to publish, or preparation of the article. Any opinions, findings and conclusions or recommendations expressed in this material are those of the author(s) and do not reflect the views of the funders.

## Supplemental Data

**Supplemental Method 1** Complement on vector cloning

**Supplemental Figure 1** Analyses of mosaic albino phenotype in T_1_ generation in *PDS3* mutants

**Supplemental Figure 2** Verification of transgene-free T_2_ albino plants

**Supplemental Figure 3** A-to-G conversion in *DM3* using *pECNUS1, pECNUS2* or *pECNUS3*

**Supplemental Figure 4** A-to-G conversion in *DM3* at Q171 target

**Supplemental Figure 5** Off-target detection in base-edited T_2_ lines for T359 target

**Supplemental Table 1** List of constructs used in the assembly of ABE vectors.

**Supplemental Table 2** List of primers used in this study.

